# Chromosome fusion affects genetic diversity and evolutionary turnover of functional loci, but consistently depends on chromosome size

**DOI:** 10.1101/2021.01.06.425547

**Authors:** Francesco Cicconardi, James J Lewis, Simon H Martin, Robert D. Reed, Charles G Danko, Stephen H Montgomery

## Abstract

Major changes in chromosome number and structure are linked to a series of evolutionary phenomena, including intrinsic barriers to gene flow or suppression of recombination due to chromosomal rearrangements. However, chromosome rearrangements can also affect the fundamental dynamics of molecular evolution within populations by changing relationships between linked loci and altering rates of recombination. Here, we build chromosome-level assembly *Eueides isabella* and, together with the chromosome-level assembly of *Dryas iulia*, examine the evolutionary consequences of multiple chromosome fusions in *Heliconius* butterflies. These assemblies pinpoint fusion points on 10 of the 21 autosomal chromosomes and reveal striking differences in the characteristics of fused and unfused chromosomes. The ten smallest autosomes in *D. iulia* and *E. isabella*, which have each fused to a longer chromosome in *Heliconius*, have higher repeat and GC content, and longer introns than predicted by their chromosome length. Following fusion, these characteristics change to become more in line with chromosome length. The fusions also led to reduced diversity, which likely reflects increased background selection and selection against introgression between diverging populations, following a reduction in per-base recombination rate. We further show that chromosome size and fusion impact turnover rates of functional loci at a macroevolutionary scale. Together these results provide further evidence that chromosome fusion in *Heliconius* likely had dramatic effects on population level processes shaping rates of neutral and adaptive divergence. These effects may have impacted patterns of diversification in *Heliconius*, a classic example of an adaptive radiation.

## Introduction

Structural changes in the genome can be an important factor for speciation (1), population divergence, and adaptation (2). Although studies of structural evolution often focus on small to medium-scale structural variants, such as inversions or translocations, large-scale structural changes, like chromosome fusion, can have substantial and immediate evolutionary impacts. The consequences of chromosome fusions occur via two main effects. First, fusions can disrupt meiotic sorting of chromosomes in heterozygous individuals or result in imbalanced gametes, both of which can lead to reproductively isolated “chromosomal races” (3–5). Second, fusion events reduce the number of unlinked DNA molecules, which results in less independence among loci. This lower per chromosome recombination rate may be favored in certain circumstances. For example, fusions may aid adaptation if recombination is reduced between co-adapted alleles at multiple loci (6). A beneficial alteration of the recombination landscape might explain why chromosomal fusions could become fixed despite initial deleterious impacts on meiosis in species with monocentric chromosomes (7, 8).

The altered recombination rate associated with chromosome fusions can have permanent downstream consequences, affecting both the efficacy of purifying selection (9) and the impact of selection on diversity at linked sites, which consequently determine levels of genetic diversity (10, 11). Indeed, although the effect is modest, genome-wide nucleotide diversity in 38 butterfly species was found to be correlated with chromosome number (12). While neutral DNA is highly susceptible to non-selective evolutionary processes such as variation in recombination rate, functional DNA elements (*e.g*., genes, enhancers, promoters, etc.) may be more constrained due to selective forces. Turnover of functional DNA (e.g., gain or loss of a CRE) is strongly associated with divergence time between taxa (13–15), but also mechanisms and processes that implicate selection, such as epistasis, gene pleiotropy, adaptive population divergence, and positive selection from environmental pressures (15–20). Thus, the extent to which changes in recombination rate might alter functional DNA evolution remain relatively unknown, and prior studies seemingly support the potential for both strong and weak effects.

In *Heliconius* butterflies, 20 of the 31 ancestral chromosomes are thought to have fused, resulting in 21 chromosomes, with the ancestral chromsome number still present in their sister genus *Eueides* (21). The impact of chromosome fusions on the recombination landscape may be especially pronounced in *Heliconius*, as holocentric species may escape some of the deleterious effects of chromosome fusions (22) and there appears to be a strict rule of one crossover per chromosome per meiosis (21). Fused chromosomes should therefore have a reduced recombination rate per base pair (bp) compared to their ancestral, unfused progenitors. In *H. melpomene*, genetic diversity is strongly negatively correlated with chromosome length (23), and longer chromosomes show reduced levels of introgression between *Heliconius* species (24, 25). Therefore, the fusion events that produced the ten longest chromosomes

## Results & discussion

### *Chromosome level assembly of the* Eueides isabella *genome and serial chromosome fusion at the origin of* Heliconius

Using a combination of high coverage, long-read (Pacific Biosciences reads) and short read data, we generated a highly complete de novo assembly *E. isabella* (Figure S1; Supplementary Material online). Final chromosome-level scaffolding brought the contiguity to 14.7 Mb with a final genome size of ~440 Mb, and very minimal N/X characters (<55 kb) (Figure S2-S4). Genome sequence composition was highly similar between *E. isabella*, *D. iulia*, and the *H. melpomene* and *H. erato* reference genomes (Table S1). Genome completeness, as measured using the BUSCO database (insect_db9: 1658 gene), is high, with 99.3%, of genes being complete (single + duplicated) and only 0.5 % (9/1658) of genes missing. Again, extremely similar results were found for *D. iulia* and the two *Heliconius* genomes (Table S1 and Figure S5). Therefore, these species represent four of the highest quality lepidopteran genomes with which to answer our questions.

In the *E. isabella* assembly the expected 31 chromosomes were recovered in 31 putative sequences, showing a highly conserved synteny with *D. iulia*, with few putative inversions (mostly at the end of chromosomes) or translocations (Figure 1, S6-S8). This allowed us to identify the ancestral-chromosome fusion points in the two *Heliconius* species with a high degree of accuracy, all of which were close to their previously estimated locations (21). We also combined a range of annotation methods to predict 26,555 protein-coding genes in the *E. isabella* genome, (Table S1). This estimate is slightly higher than the other Heliconiini genomes, probably due to a higher degree of gene model fission as indicated by the lower distribution of completeness (Figure S9). Repetitive elements account for 29.80% of the *E. isabella* genome, which is slightly less than in *D. iulia* and *H. erato* (31.74% and 33.02%, respectively), but significantly more than *H. melpomene* (25.90%; Table S1; Figure S10). In keeping with other Heliconiini genomes, total interspersed repeats constitute the largest fraction of repetitive elements (23.91%).

**Figure 1.**
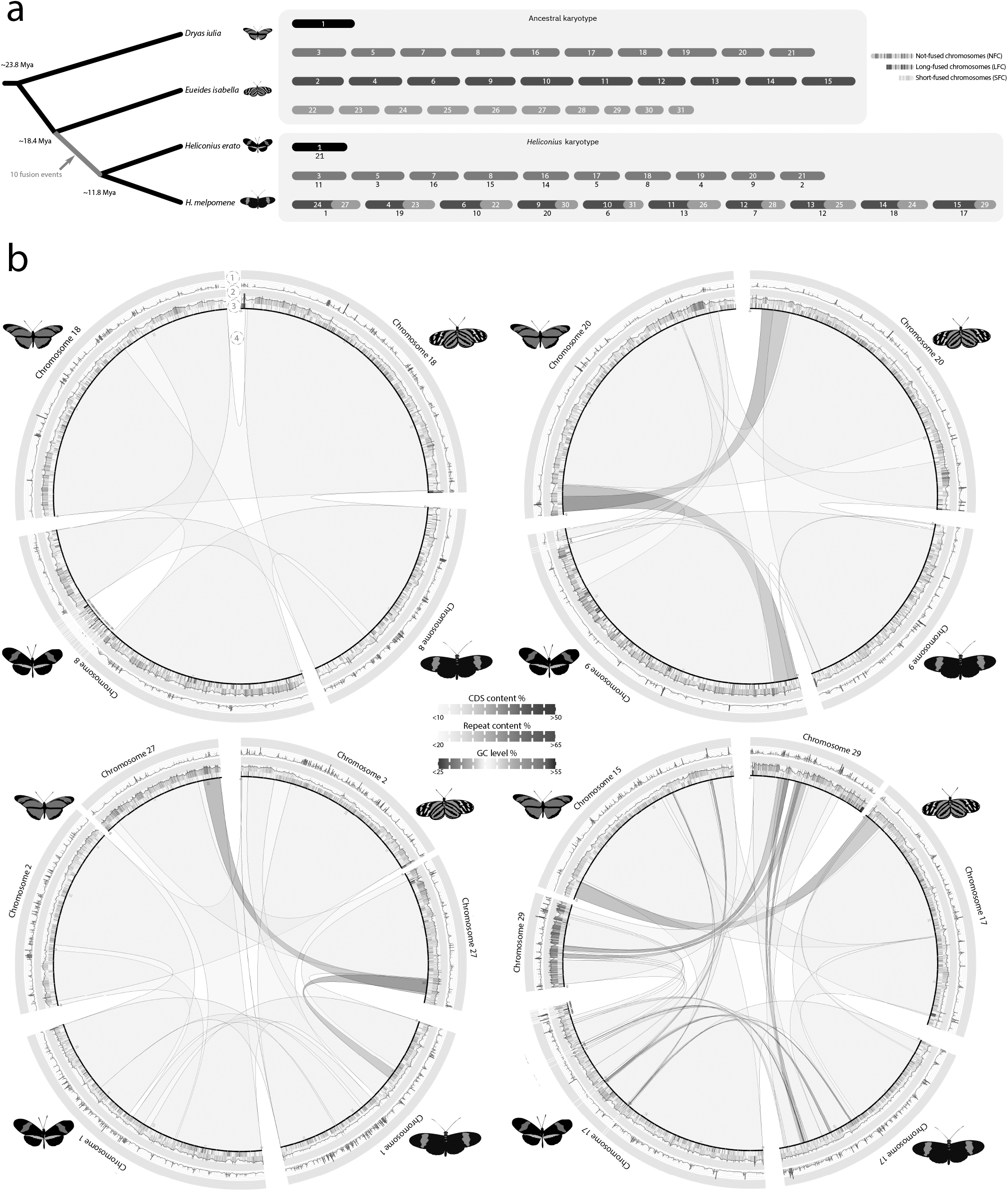
Synteny plots for four representative chromosome fusions. The comparisons are shown only for the flanking species, in order to avoid confusion. For layer of for the plots indicate: 1) Coding region content, higher density is shown in darker red; 2) Repetitive element richness, higher density is shown in darker blue; 3) GC level, in red AT-rich regions while in blue more GC-rich content; 4) Ribbons showing synteny blocks (light blue), blocks are red colored when the block is inverted in the comparison. The ideogram scale is in Mbp.

### Chromosome fusion leads to a change in the spatial distribution of repetitive elements

Comparative analyses of chromosome composition in lepidoptera has shown that the distribution of repeats, GC and gene content don’t follow a consistent pattern across chromosome length (28). We explored these features in our chromosome level assemblies, looking for an effect of chromosome fusions on chromosome composition. We found that across all four species, chromosome lengths are highly correlated with repeat content (*P* < 0.0001; Pearson’s *ρ* = 0.89; Figure S11). All chromosomes in *D. iulia*, *E. isabella*, and the unfused chromosomes in in *Heliconiu*s likely altered the evolutionary trajectories of the ten pairs of shorter progenitor chromosomes involved. Indeed, Davey et al. (21) hypothesised that the altered genome structure may contribute to the elevated speciation rate in *Heliconius* (26). Here, we aimed to test whether increases in chromosome length have altered levels of nucleotide diversity and rates of evolutionary turnover in *Heliconius*. We combine our newly available *Dryas iulia* genome assembly (27), with a chromosome level assembly of *Eueides isabella*. We then compare these genomes, which have the ancestral chromosome number of 31, with the genomes of *H. melpomene* and *H. erato*. Our findings show that the ten smallest chromosomes in *Dryas* and *Euiedes*, which are fused in *Heliconius*, have a distinct genomic and gene-structural composition. Chromosome fusions are associated with a dramatic change in recombination and background selection. Finally, we show that at a macroevolutionary time scale this had a significant effect on DNA content putatively under selection.

*Heliconius* have higher GC and repetitive element frequencies toward the chromosome ends. In contrast, coding regions have slightly higher frequencies in the middle of the chromosomes (Figure 2a-b). This pattern is maintained among fused *Heliconius* chromosomes and their unfused homologous chromosomes in *D. iulia* and *E. isabella*. If CG-rich and repetitive element accumulations in the chromosome’s tails were retained after chromosome fusions, then fusion events should produce a W-shaped distribution centered around the fusion regions; however, no trace of this was found (Figure 2b). This suggests selection acted to remove repetitive elements in the central part of chromosomes. Comparing the ten smallest Heliconiini chromosomes with their homologues in *Melitaea cinxia,* it seems clear that chromosome lengths have been highly stable over this long evolutionary time period. Our finding seems to contradict the notion that holocentric chromosomes have uniform distributions of GC, repeats and gene content across chromosomes (22, 29), contrasting with monocentric chromosomes with localized centromeres, in which they are compartmentalized to GC-rich and GC-poor regions (30). Whether the uneven distributions we observe are favoured due to meiotic pairing or some other selective driver remains to be tested.

**Figure 2.**
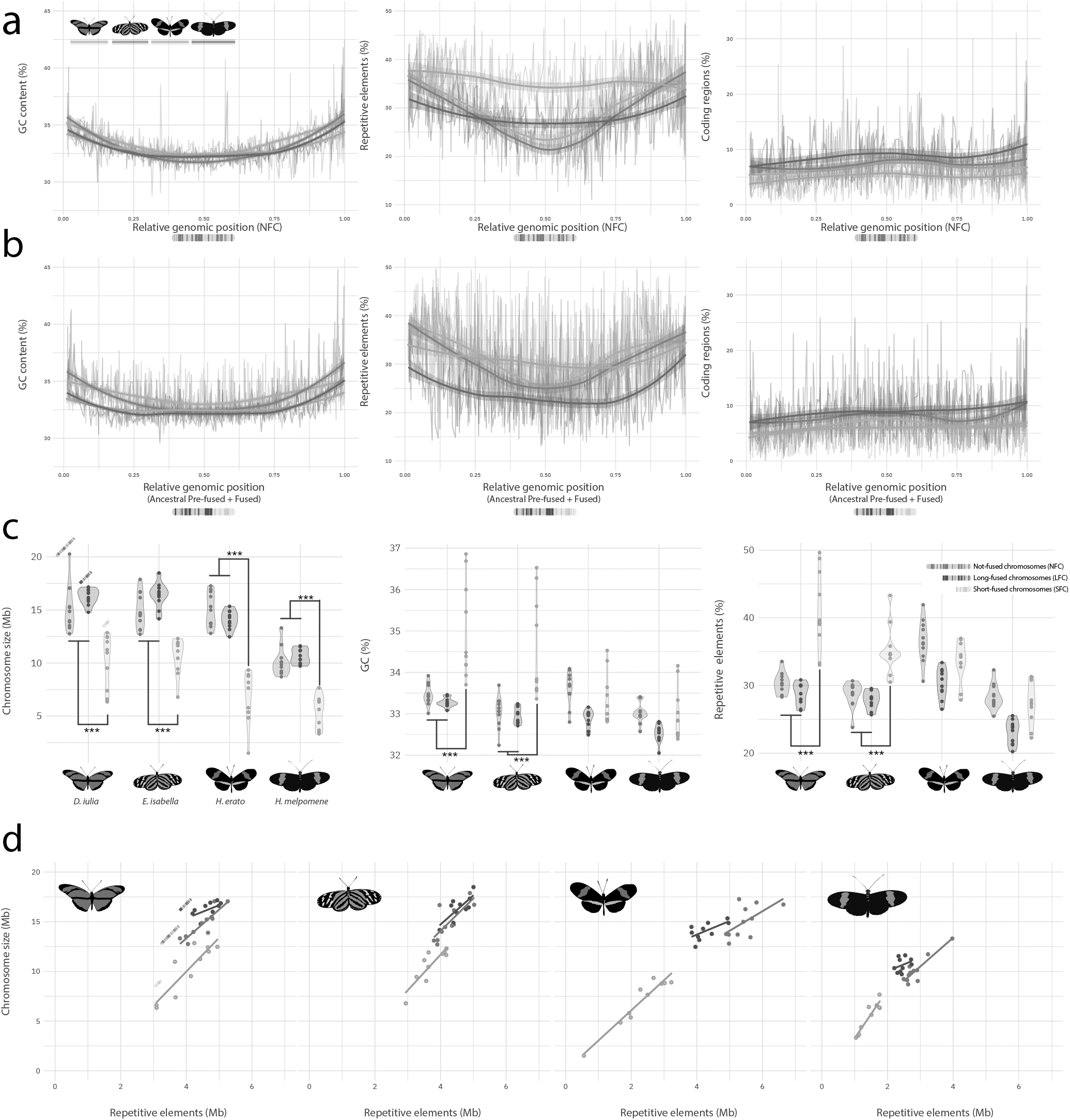
a) GC content (%), repeat content (%) and coding density within 100 kb non-overlapping windows in not fused chromosomes (NFC) of the four species (showed in colour). b) same as a) but for Heliconius fused chromosomes and their homologous in *D. iulia* and *E. Isabella*. In both cases, GC and repeat distributions are accumulating at the chromosome tails, while coding density seems to be slightly denser in the central part of chromosomes. c) Violin plots of the different chromosome types, showing how and where size, CG content (%) and repetitive elements (%) are different. Short-fused chromosomes (SFC) appear to be smaller and with higher GC and repeat content compared with not-fused (NFC) and long-fused (LFC) chromosomes, which, in turn, to not appear significantly different. *** Wilcoxon rank-sum test ‘less’; *P* values less than 0.001. d) Scatter plots between chromosome size and their repetitive element (Mb), their relative regression line, for the different chromosome types (colour) for each species.

### *Chromosomes that were fused in* Heliconius *are small and have distinct nucleotide compositions*

Previous comparisons between *Bombyx mori, M. cinxia,* and *H. melpomene* suggested that lepidopteran chromosomal fusion events involved the shortest chromosomes (28). These tend to have high rates of chromosomal evolution and rearrangement and high proportions of repetitive elements (see above). These factors might explain the disproportionate involvement of short chromosomes in fusions in *Heliconius* (28). The accurate chromosome level-assemblies of *E. isabella* and *D. iulia* allowed us to test whether this pattern is consistent in a shallower phylogenetic framework, identifying how chromosome fusion events have shaped *Heliconius* genomes with greater precision.

To do so we split the fused chromosomes in *Heliconius* into their corresponding pre-fused homologous chromosomes. These were categorized as: i) not-fused chromosomes (NFC), homologous chromosomes that are conserved within Heliconiinae; ii) long-fused chromosomes (LFC), the longer split chromosome sequence in *Heliconius* and its ancestral homolog; and iii) short-fused chromosomes (SFC), the shorter split chromosome sequence in *Heliconius* and its ancestral homolog. Across all species, SFCs are the smallest chromosomes (Wilcoxon rank-sum test ‘less’; *P* ≤ 4.5×10^−08^), even in *D. iulia* and *E. isabella* (Wilcoxon *P* < 0.01). *H. melpomene* has smaller chromosomes overall (22) compared to the two outgroups (Wilcoxon *P* ≤ 1.3×10^−11^), but in *H. erato* only the fused chromosomes, both LFC and SFC, differ in size from *D. iulia* and *E. isabella* (Figure 2c-d; Wilcoxon *P* ≤ 0.00016). This indicates that *H. melpomene* experienced a secondary reduction in genome size after the fusion events with reductions in chromosome size being specific to the fused chromosomes, while *H. erato* did not (Figure S12).

Within *D. iulia* and *E. isabella*, pre-fused SFCs displayed a different composition from other chromosomes, with both GC content and repetitive elements being higher in SFCs (Wilcoxon rank-sum test ‘greater’; *P* ≤ 6.7×10^−06^) (Figure 2c). The relationship between chromosome size and repetitive elements other Heliconiini: *P*-adjusted ≤ 0.0018), followed by *E. isabella* and *H. erato* (0.31±0.10 and 0.39±0.07, respectively) and finally *H. melpomene* (0.53±0.04; *P*-adjusted ≤ 8.2^−06^) (Figure 2d; Table S2). Hence, the more typical composition of across species and chromosome types (NFC, LFC and SFC) also shows striking differences. Our analysis revealed significant shifts in the distribution of repetitive elements, relative to other sequence elements, between our three chromosome categories (Figure S11). This indicates that rates of repetitive element expansion/contraction vary predictably between chromosome types. SFCs show particularly high interspecific variation in chromosome composition, with *D. iulia* having the lowest repeat content, for a given chromosome length (intercept = 0.23±0.11; vs. SFCs in *Heliconius* is explained by SFCs shifting in composition to match those of larger chromosomes, the LFCs and NFCs.

As recently reported in millipedes (31), genomes with more repetitive elements show significant expansions in intron size. We explored this effect in *Heliconius* by looking at the composition of annotated gene models in SFCs. While transcript, CDS, exon, and UTR lengths show highly similar distributions across species, intron lengths differ between both species and chromosome types. *D. iulia* has significantly longer intron sizes (Wilcoxon rank-sum test ‘greater’; *P* < 2.2×10^−16^; mean: 2688 bp), followed by *E. isabella* (Wilcoxon rank-sum test ‘greater’; *P* < 2.2×10^−16^; mean: 2335 bp)*, H. erato* (Wilcoxon rank-sum test ‘greater’; *P* < 2.2×10^−16^; mean: 1770 bp) and *H. melpomene* (mean: 972 bp) (Figure S14). We further found that, within species, intron length for genes on SFCs are significantly longer than for LFCs or NFCs (Wilcoxon rank-sum test ‘greater’; *P* ≤ 2.63×10^−14^) (Figure S15). This pattern is highly similar to that of repeat content suggesting a correlation between the two (31). Indeed, SFC introns contain significantly higher amounts of transposable elements than would be predicted by their length (*P* ≤ 6.7×10^−09^; elevation for intergenic regions 0.79±0.03; elevation for intronic regions 0.67±0.02). In contrast, repeat content for introns and intergenic regions was not significantly different in NFCs and LFCs. For *D. iulia* and *E. isabella* the regression slopes, in terms of allometric differences, showed no significant difference (*P* ≥ 0.15), but elevation along the *y*-axis did, with intronic elevation ~50% lower than intergenic regions (*P* < 2.2×10^−16^; Table S4), reflecting with the higher amount of repetitive elements in *Dryas*. Overall, these findings show how repetitive element content influence not only genome size but also the gene structure itself. Why introns are more affected by repetitive elements is not clear. It is possible that purifying selection may be stronger on intergenic regions, perhaps to avoid the disruption of regulatory elements in relatively compact genomes. Alternatively, the presence of transposable elements in intronic regions may favor exonization, the acquisition of new exons from non-protein-coding, primarily intronic, DNA sequences (32).

### Chromosome fusions caused reductions in recombination rate and levels of diversity

Although the fused chromosomes in *Heliconius* tend to be much longer than the unfused chromosomes (by ~40-60% in *H. erato* and *H. melpomene* respectively), genetic linkage maps (20, 21) reveal that map lengths are consistently close to 50 centimorgan (cM) for all chromosomes, with fused chromosome maps only marginally longer than unfused (by 5% in *H. melpomene* and 7% in *H. erato*) (Figure S17). This implies an average of one crossover per pair of bivalents per meiosis, regardless of chromosome length (21). If applied across the 31 ancestral chromosomes, this would imply that fused chromosomes, especially SFCs, have experienced a dramatic decrease in their per-base recombination rate. For example, ancestral chromosome 31, an SFC forming part of chromosome 6 in *Heliconius*, accounts for just 6 cM of the total 48 cM (Figure S17). We therefore hypothesized that the decrease in recombination rate would result in a decrease in genetic diversity for the SFCs and LFSc, as recombination rate determines the extent to which the processes of background selection and genetic hitchhiking tend to reduce diversity at linked sites (10, 33).

As a proxy for the level of neutral genetic diversity in each of the four species, we computed nucleotide diversity (π) at 4-fold degenerate 3rd codon positions (hereafter π4D) using re-sequenced individuals (see Table S5 for sample information and accession numbers). There is a strong negative relationship between chromosome length and π4D in *D. iulia* (*R*^*2*^ = 0.506, *P* = 1×10^−5^), *H. melpomene* (*R*^*2*^ = 0.752, *P* = 1×10^−6^) and *H. erato* (*R*^*2*^ = 0.894, *P* = 3×10^−10^) (Figure S18), consistent with lower per-base recombination rates on longer chromosomes resulting in greater levels of background selection and/or hitchhiking (23). This trend was not seen in *E. isabella*, which generally shows low genetic diversity across all chromosomes. It is possible that a reduction in effective population size has reduced the efficacy of background selection in *E. isabella* such that chromosome-level recombination rate is no longer a good predictor of diversity. We therefore focused on *D. iulia* as representative of the ancestral state for comparison with the *Heliconius* species.

Relative levels of diversity tend to be reduced on both the LFCs and SFCs in *Heliconius* compared to their unfused homologues in *D. iulia* (Figure 3a,b). The difference is most pronounced for the SFCs, which tend to have experienced the greatest decrease in recombination rate following the fusion events. Indeed, assuming map lengths of 50 cM for *D. iulia* chromosomes, we can estimate recombination rates for each chromosome before and after the fusions. This estimated reduction in recombination rate is a strong predictor of the reduction in relative levels of diversity (Spearman’s *ρ* = 0.757, *P* = 3×10^−6^ fo *H. melpomene*; *ρ* = 0.662, *P* = 1×10^−4^ for *H. erato*; Figure 3c,d).

**Figure 3.**
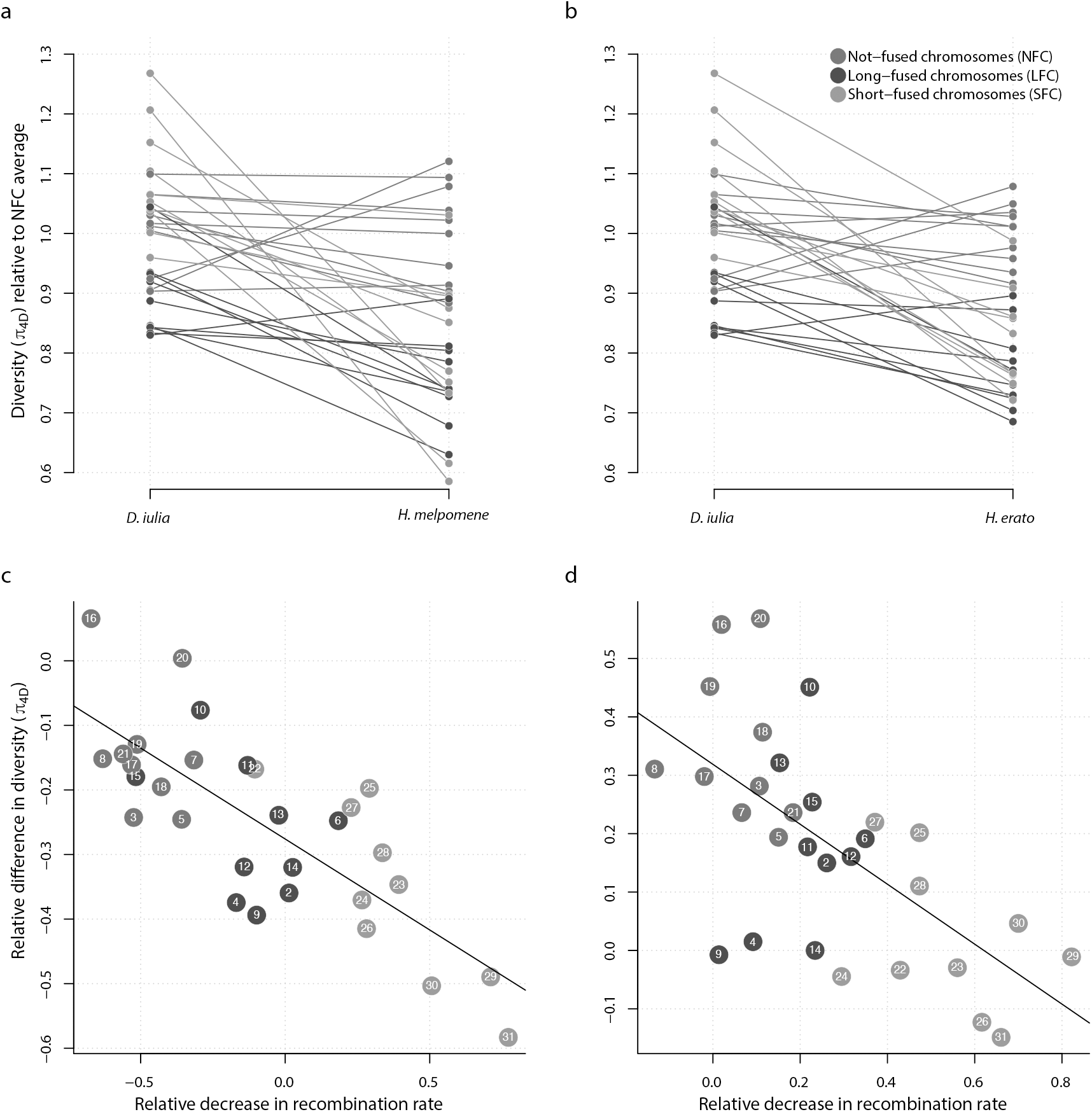
Loss of genetic diversity following chromosome fusions in *Heliconius*. (a, b) Nucleotide diversity at four-fold degenerate codon positions (π4D) averaged across each of the 31 ancestral chromosomes, and normalised relative to the average π4D for not-fused chromosomes (NFC) to account for different effective population sizes between species. The outgroup species *D. Iulia* is compared to *H. melpomene* (a) and *H. Erato* (b). Downward sloping lines indicate chromosmes that have lower relative genetic diversity in Heliconius. (c, d) The relative change in π4D plotted as a function of the relative decrease in recombination rate following the 10 fusion events in in *H. melpomene* (c) and *H. erato* (d). We assume that the ancestral chromosomes had lengths and levels of diversity equal to those in *D. iulia*, and recombination map lengths of 50 cM each (see main text). Points further to the right indicate chromosomes that have experienced a greater decrease in recombination rate. Note that in *H. melpomene* the NFCs show negative values on the x-axis, indicating an increase in recombination rate, due to the genome-size reduction in *H. melpomene*. Nonetheless, the overall trend that the decrease in relative recombination rate has been greatest for fused chromosomes (especially SFCs) is present in both H. melpomene and H. erato. A fitted linear regression line is included for convenience.

Our findings demonstrate that the ten fusion events in the ancestor of all *Heliconius* species resulted in a dramatic change in the fate of the chromosomes involved. First, the reduced effective population size caused by increased background selection likely also leads to a reduction in the efficacy of selection. Adaptive evolution may be further impeded by Hill-Robertson interference between tightly linked selected loci (34, 35). Evidence for less efficient selection in genomic regions of reduced recombination rate has been reported in *Drosophila* (36–38). Given that the gene complement of each fused chromosome in *Heliconius* is largely unchanged from that in outgroup species, we hypothesise that the fusions may have resulted in a systematic shift in the relative importance of certain genes for adaptation. A second side-effect of reduced recombination rate in the fused *Heliconius* chromosomes is an increase in barriers against introgression between hybridising species (24, 25). This occurs because introgressed tracts take longer to break down in regions of lower recombination rate and are consequently more rapidly purged by selection. This raises the intriguing possibility that the fusion events caused some regions of the genome to become less permeable to gene flow (21). Future work on these, and other, taxa will further detail the importance of chromosome fusions (and fissions) in determining the subsequent evolutionary trajectory of different genomic regions.

### Chromosome fusions alter the rate of turnover in functional DNA

Finally, we sought to assess how the ten chromosome fusions in *Heliconius* have affected loci putatively under selection. While it is clear that chromosome fusions can drive a decrease in neutral nucleotide diversity, the extent to which the recombination landscape alters the evolution of functional elements in the genome remains uncertain. Functional elements, such as protein coding sequences and cis-regulatory elements, likely face opposing forces from neutral processes driven by reduced recombination rates and purifying or directional selection (Villar 2015, Lewis 2016, Lewis 2019 MBE). We therefore assessed evolutionary turnover of accessible chromatin and protein coding exons to test whether the fusion of *Heliconius* chromosomes had a similar effect on functional loci as we observed at neutral sites. We used DNA sequence conservation of cis-regulatory elements (CREs) and gene exons from *D. iulia* to set a baseline for functional DNA turnover in the Heliconiini lineage. As expected, divergence time strongly effected conservation of both functional categories. We observed a similar degree of conservation between annotated *D. iulia* functional DNA and the *E. isabella* and *Heliconius* genomes, which represent 26 million years of sequence divergence (Figure 4a). Conservation continued to decrease with divergence time in comparisons against *M. cinxia* (65 million years diverged) and *Danaus plexippus* (85 million years diverged). For each species comparison, however, both exons and CRE DNAs conservation displayed a strong correlation with chromosome length (Pearson’s *ρ* = 0.42-0.78). The correlation between functional DNA conservation and chromosome length decayed slightly with increased divergence time, most notably in exons, which showed a higher overall correlation (e.g., *ρ* = 0.078 for *H. melpomone* DNA) at lower divergence times. Thus, while some additional forces may affect functional DNA turnover, these results confirm that functional DNA elements remain subject to the same effects of recombination rate as neutral loci over macroevolutionary time.

**Figure 4.**
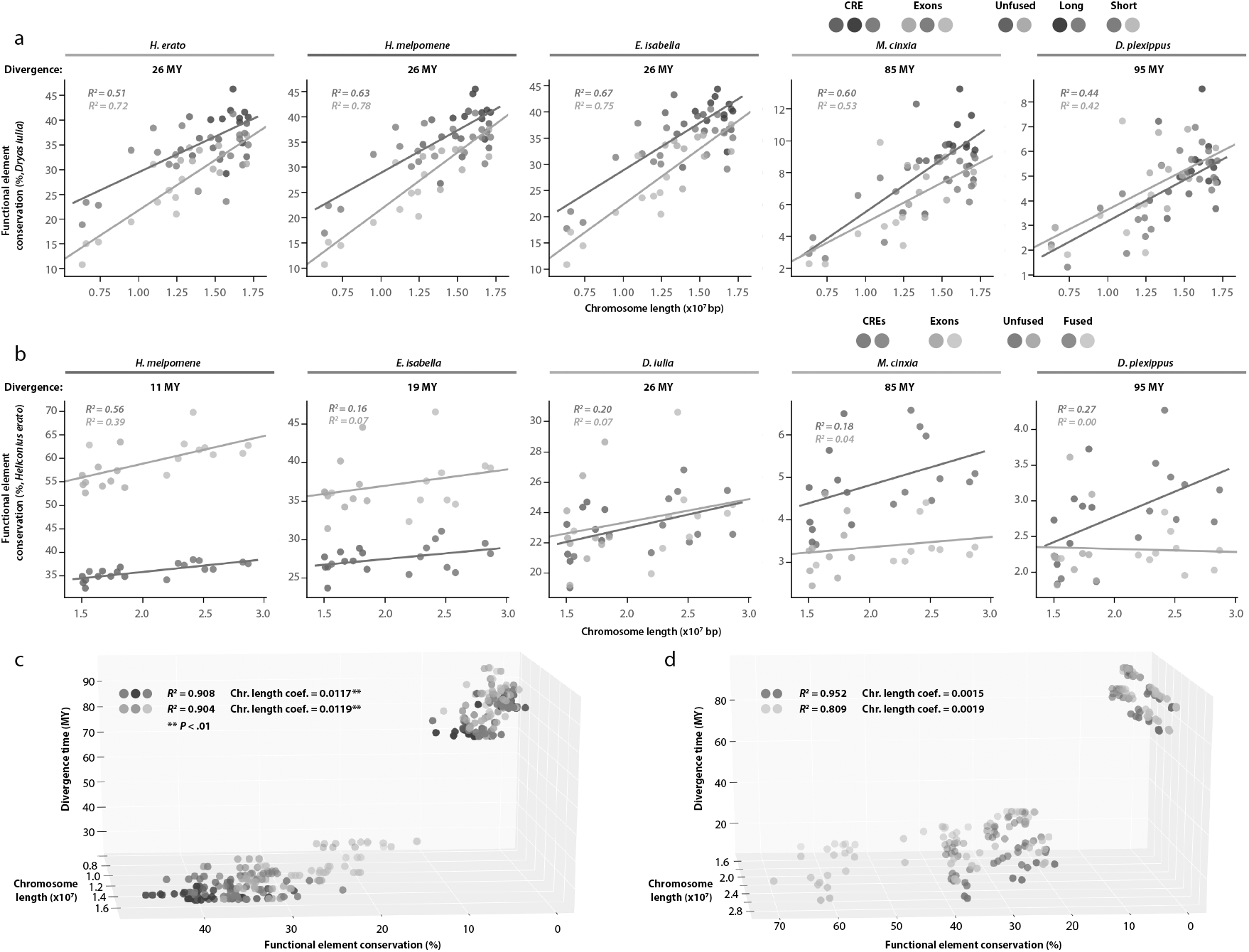
Patterns of functional divergence in fused and unfused chromosomes. (a) Percent of *D. iulia* CREs (dark circles) and gene exons (light circles) conserved over 95 million years of divergence. Long (purple), short (brown?), and unfused (green) chromosomes all consistently show a strong correlation of DNA conservation with chromosome length. (b) Percent of *H. erato* CREs (dark circles) and gene exons (light circles) conserved over 95 million years of divergence. Chromosome length is highly correlated with functional DNA conservation between *H. erato* and *H. melpomene*, both of which have 21 chromosomes. This correlation declines rapidly with divergence time and comparison with unfused chromosomes. Multiple regression analysis of chromosome length and divergence time on functional DNA conservations shows that chromosome length is significantly more impactful on *D. iulia* sequences (c) than for H. erato CRES or gene exons (d).

To determine the extent to which chromosome fusions, and change in recombination rate, may have altered the turnover of functional DNA in *Heliconius*, we performed the same DNA conservation analysis using CREs and exons from *H. erato*. Testing the complete *H. erato* chromosomes allowed us to determine the effect of changes in recombination rate on functional DNA content over the past 11-19 million years post-fusions. In contrast to our analysis of *D. iulia* functional DNA, *H. erato* showed a fairly strong correlation with *H. melpomene* (Exons: *ρ* = 0.39, CREs: *ρ* = 0.56), but a substantial loss of association between chromosome length and DNA conservation between *H. erato* and more distantly related species with 31 chromosomes (Exons: *ρ* = 0.07, CREs: *ρ* = 0.16).

The reduced correlation between chromosome length and functional DNA conservation suggests that the relatively recent change in recombination rate in *Heliconius* may not have impacted functional DNA turnover to the same extent as stable recombination rates over a much longer period of time. To test this more explicitly, we performed multiple linear regression of *D. iulia* (Figure 4c) and *H. erato* (Figure 4d) functional elements to determine the effect of chromosome length relative to divergence time. Multiple regression models for both *D. iulia* and *H. erato* functional elements accounted for the substantial majority of variance in functional DNA conservation (*r*^2^ = 0.81-0.95). There was, however, a marked and significant (paired *t*-test, *P* < 0.01) difference in the effect size of chromosome length between regression models of *D. iulia* and *H. erato* functional DNA. The impact of chromosome length and, by extension, recombination rate on DNA conservation was approximately seven times greater in *D. iulia* than for *H. erato*. The contrast between *D. iulia* and *H. erato* exon and CRE turnover thus suggests that recombination rate has a significant effect on DNA content putatively under selection, but that changes in recombination rate are likely more impactful on average rates of change across chromosomes over long periods of macroevolution than over shorter divergence times within genera. This impact may not be homogenous across the chromosome, however, and thus local changes in recombination rate may be important over microevolutionary timescales.

## Conclusions

Our analyses of four high-quality chromosome-level assemblies *D. iulia*, *E. isabella, H. erato* and *H. melpomene* reveal important insights into the evolutionary impact of chromosome fusion. We have shown that Nymphalidae chromosomes differ significantly by size, both in terms of nucleotide composition and gene structure. Specifically, chromosome length is inversely correlated with repetitive element content possibly making them more likely to fuse with longer, more stable chromosomes. Interestingly, both whole genomes and indivdiual chromosomes with higher repetitive element content have longer introns on average, which is due to a higher denisty of repeats compared to intergenic regions, a feature that in theory could lead to the formation of new functional exons. We further show that the ten fusion events resulted in a dramatic change in the fate of the chromosomes involved, reducing the effective population size by increasing background selection, possibly resulting in a systematic shift in genes targeted divergent selection and barriers to gene flow. Finally, we explored if the chromosome fusions may have also changed rates of evolutionary turnover of accessible chromatin and protein coding exons and found that both exons and CREs conservation displayed a strong correlation with chromosome length, confirming that functional DNA elements are subject to the same effects of recombination rate over macroevolutionary time. These findings, and the genomic resources we provide, are not only important to understand how genomic architecture impacted lepidopteran evolution, but also to understand how the karyotype can affect species evolution at a broader biological scale.

## Acknowledgements

We thank the supported by NASA 17-EXO-17-2-0112 to CGD, NSF IOS-1656514 and DEB-1546049 to RDR, and a NERC IRF (NE/N014936/1) and ERC Starter Grant (758508) to SHM SM was supported by the Royal Society URF\R1\180682. FC would like to thank Dr. Pasi Rastas for his help in reconstruct the linkage map and Angel Corpuz for various informatics support. FC and SHM are grateful to the High-Performance Computing team at the Advanced Computing Research Centre, University of Bristol for support.

Funders, AHRC people, Pasi, Gepi

## Methods

### DNA and RNA extraction and sequencing

Individuals of *E. isabella* were collected from partially inbred commercial stocks (Costa Rica Entomological Supplies, Alajuela, Costa Rica). High-quality, high-molecular-weight genomic DNA was extracted from late stages of pupae, dissecting up 100mg of tissue, mainly thorax and wing imaginal disk, frozen in liquid nitrogen and homogenized in 9.2ml buffer G2 (Qiagen Midi Prep Kit) adding 19 μl of RNAseA. The samples were then transferred in a 15ml tube adding 0.2 μl of Protease K and incubated at ~50 °C for two hours. Samples were transferred to Genomic tip and processed with Qiagen Midi Prep Kit (Qiagen, Valencia, CA) following the manufacturer’s instructions. DNA was then precipitated using 2ml 70% EtOH and dissolved in water.

From the same stocks, RNA was extracted separately from six adult tissue (four wings; three heads; four antennae, legs and mouth parts; thorax; abdomen 1-3, abdomen 4-6), and five tissue parts from early ommochrome stage of the pupae (head and mouth parts; wings, antennae and legs; thorax; abdomen 1-3, abdomen 4-6). Each tissue was frozen in liquid nitrogen and quickly homogenized in 500 μl Trizol, adding the remaining 500 μl Trizol at the end of the homogenization. Phase separation was performed adding 200 μl of cold chloroform. The upper phase was then transfer to RNeasy Mini spin column and processed with Qiagen RNeasy Mini Prep Kit (Qiagen, Valencia, CA) and DNAse purification using the Turbo DNA-free kit (Life Technologies, Carlsbad, CA, USA) following the manufacturer’s instructions. All the extractions were finally pooled keeping the same final RNA concentration from all samples.

The Pacific Bioscience (PacBio) data were then sequenced at the Centre for Genomic Research, University of Liverpool using PacBio sequel SMRT cell (2.0 chemistry), while polyadenilated Illumina RNAseq data (125bp x 2) where generated at the Institute of Applied Genomics (IGA), Udine, Italy.

### Genome assembly

PacBio reads were corrected, trimmed and assembled using CANU v.1.8 +356 changes (settings: genomeSize = 400m; corMhapSensitivity = normal; corOutCoverage = 100; correctedErrorRate = 0.105; ovlMerThreshold = 500; batOptions = -dg 3 -db 3 -dr 1 -ca 500 -cp 50) (39). One problem with this type of long-read sequencing technology is the presence of post-assembly insertions and deletions (indels) (40). CANU can correct reads but does not remove all indels. The resulting raw assemblies were therefore corrected by remapping all uncorrected raw PacBio reads with pbmm2 v.1.0.0 and correcting the assembly with Arrow v.2.3.3 (https://github.com/PacificBiosciences/GenomicConsensus) for three iterations. To further error correct for indels we used short Illumina reads for *E. isabella* (41) with Pilon v.1.23 (42) for five iterations. Assemblies were then processed with Purge Haplotigs v.20191008 (purge -a 85) (43) in order to remove haplocontigs, artificially duplicated genomic regions due to heterozygosity. To correct for mis-assemblies Polar Star (https://github.com/phasegenomics/polar_star) was employed. This algorithm calculates read depth of aligned PacBio reads to the assembly at each base. Read depth is then smoothed in a 100 bp sliding window, and regions of high, low, and normal depth are merged. Low read depth outliers are identified, and contigs are broken at each such location. Contigs were then re-scaffolded using P_RNA_scaffolder (44), which uses information from RNAseq mapping, and LRScaf v.1.1.5 (45), which uses information from long-reads. The resulting gaps were then filled using PacBio reads applying LR_Gapcloser v.1.1 (46). After the introduction of this new PacBio information, we repeated the previous polishing procedure using five iterations of Pilon, plus three more iterations with Illumina RNA-seq data to correct indels only. Before the chromosome level scaffolding, we used synteny maps implemented with BLAST (47) and ALLMAPS (48) to identify duplicated regions at the end of scaffolds, trimming them away.

Illumina RAD-seq data from Davey et al. (2016) were downloaded and aligned to the scaffolds of *E. isabella* assembly using STAR v.2.7.2c (49) to generate the linkage map for the final scaffolding step. Duplicate reads were removed using the Picard v.2.0.1 MarkDuplicates tool (http://broadinstitute.github.io/picard). Indels were subsequently realigned using the RealignerTargetCreator and IndelRealigner tools from GATK v.3.8-1-0-gf15c1c3ef (50). Individual SNPs were called using mpileup in SAMtools (51) in combination with pileupParser2 and pileup2posterior modules from Lep-MAP v.3 (52). SNPs were filtered using Lep-MAP Filtering2 and the final linkage generated using SeparateChromosomes2, JoinSingles2All and OrderMarkers2 (parameters: dataTolerance = 0.001; distortionLod = 1; lodLimit = 14; femaleTheta = 0; maleTheta = 0.05). The scaffolding stage was performed with Lep-Anchor (52).

### Transcriptome annotation

RNA-seq raw read data were filtered using Trimmomatic v.0.39 (53) (ILLUMINACLIP:$ILLUMINACLIP:2:30:10; SLIDINGWINDOW:5:10; MINLEN:100). RNA-Seq data were used as training data input to *ab initio* and *de novo* predictors and for direct alignments, and were mapped to the genome using STAR v.2.7.2c aligner (alignIntronMax = 500000; alignSJoverhangMin = 10). The resultant sorted BAM files were used as training for the BRAKER v.2.1.5 pipeline (54, 55), which combines GeneMark-ES Suite v.4.30 (56) and AUGUSTUS v.3.3.3 (57), along with the masked genomes, generated with RepeatMasker v.4.1.0 (58) using the Lepidoptera database and protein alignments from closely related, or model species (*Drosophila melanogaster, Bombyx mori, Bicyclus anynana, Danaus plexippus, Heliconius erato,* and *H. melpomene*) using Exonerate v.2.4.0 (59). BRAKER is an automated pipeline to predict genes that uses iterative training of AUGUSTUS to generate initial gene predictions. GeneMark-predicted genes are filtered and provided for AUGUSTUS training, followed by AUGUSTUS prediction, integrating the RNA-Seq and protein alignment information, to generate the final gene predictions.

To generate the de novo transcriptome assemblies, quality filtered reads assembled using Trinity v.2.10.0 (60) and generated transcript were subsequently aligned to the genome using Minimap2 v.2.17-r974-dirty (61). To generate the ab initio transcriptomes, reads were re-aligned to the genome using STAR with the same parameters with Braker predicted splice sites, and assembled using Stringtie v.2.1.3b (62) and Cufflinks v.2.2.1 (63, 64) for each sample. The assemblies derived from all samples within each program were merged using Stringtie --merge. All such generated transcriptomes (Trinity, BRAKER, StringTie and Cufflinks) were merged used with STAR to re-map once again all raw reads in order to evaluate all detectable splice-sites with Portcullis v.1.1.2 (65). Transcripts from ab initio and de novo assemblies (Trinity, StringTie and Cufflinks) without supported splice-junctions were therefore filtered out, and only transcripts with unique intron chain were retained from all annotations.

The best putative protein-coding sequences were finally inferred using TransDecoder v5.5.0 (http://transdecoder.github.io/) (minimum amino-acid length > 50) using homologs from UniProt database (66) and Lepidoptera proteome (see below) found with deltaBLAST v.2.7.1+ (67); and PFam domains (68) with HMMscan v.3.3 (69) (e <1e-10). For transcripts without a putative protein-coding region CPC2 (70) was adopted to identify putative non-coding transcripts.

### Chromosome analysis

Chromosome mapping was carried out using orthologous genes between *D. iulia, E. Isabella, H. erato and H. melpomene* to define the level of gene conservation and translocations among chromosomes. Synteny maps were implemented with BLAST and ALLMAPS using Minimap2 to map *H. erato* and *H. melpomene* loci to the *D. iulia* and *E. isabell*a assemblies. This let us identify the putative junctions between the two fused chromosome pairs. The midpoints between the flanking orthologs of different homologous chromosomes were used to split the fused chromosomes in *H. erato* and *H. melpomene* in two, recreating the ancestral chromosome set. Chromosome mappings are illustrated using Circos (71).

We explored the scaling relationship between chromosome size and repetitive elements across species and chromosome types (NFC, LFC and SFC) using SMATR v.3.4-8 (72), together with a Wilcoxon-Mann-Whitney rank sum test as implemented in the R function WILCOX.TEST (http://www.r-project.org) v.3 (R Development Core Team 2017) to test for differences in the relationships between chromosome size, GC content, and repetitive element content among chromosome types (chromsomes that were fused/unfused in *Heliconius*). In both cases we adopted a stringent *P*-value threshold of 0.005 (73). We also calculated GC, repetitive element and coding-sequence (CDS) content within non-overlapping 500 kb sliding windows for the chromosomes of all the species, with the order and orientation of the chromosomes determined based on synteny to *H. melpomene*. We used the annotation and a custom python script to extract strand specific intronic regions (BED format), while to extract strand specific intergenic regions the initial annotation was split into plus and minus strands and BEDTools complement v.2.29.0 (74) used to generate intronic and intergenic regions. For both annotations BEDTools getfasta were used to extract their relative sequences analysed with RepeatMasker. Their relative scaling coefficients and intercepts were subsequently analysied with SMATR as reported above.

### Analysis of genetic diversity

To investigate whether chromosome fusions are associated with long-term shifts in levels of genetic diversity, we analysed genome resequencing data from 3-6 additional individuals of each species (see Table S1 for sample details and accession numbers). For newly sequenced individuals, DNA extraction was performed using the Qiagen DNeasy Blood and Tissue kit and libraries were prepared following a low-input Nextera-based DNA library prep protocol (skim sequencing) at the Cornell Institute of Technology Core Facility. Libraries were sequenced on a HiSeq4000. Reads were aligned to their respective reference assemblies using BWA MEM version 0.7.17 (75). Processing and sorting of SAM and BAM files was performed using SAMtools version 1.9 (51), and duplicate reads were removed using Picard MarkDuplicates version 2.21.1 (https://broadinstitute.github.io/picard/). For genotyping we targeted only coding sites, as we were specifically interested in comparing levels of diversity among four-fold degenerate 3rd codon positions, which provide the most reliable comparison of relative diversity at nearly neutrally evolving sites in the genome. We therefore first defined the set of non-overlapping coding intervals in the genome based on the annotations and then used bcftools call version 1.9 (76) with the -m option (https://samtools.github.io/bcftools/call-m.pdf) to call genotypes for this subset of sites. Only individual genotypes with read depth > 10 and genotype quality (GQ) >30 were considered, except in *D. iulia*, where lower coverage sequencing meant that these thresholds had to be decreased to 5 and 20, respectively.

Four-fold degenerate sites (‘4D sites’ hereafter) were defined as 3rd codon positions at which a substitution to any other base would not alter the encoded amino acid. This condition was evaluated by considering not only the codon sequence in the reference assembly, but also any variants observed in the re-sequenced individuals, and codons with more than one valiant site were automatically discarded, resulting in the most conservative possible set of 4D sites. These steps were implemented using a custom script. Nucleotide diversity was computed in non-overlapping windows of 500 4D sites each, using the script popgenWindows.py (https://github.com/simonhmartin/genomics_general). In *D. iulia*, because three of the re-sequenced individuals were from a small Key Largo population that has reduced diversity, we instead used pairwise diversity between one Key Largo individual and the 10X Genomics sequenced individual used for genome assembly (27).

### Analysis of functional DNA turnover between species

To determine the extent to which changes in per base pair recombination rate may have alter functional DNA turnover between species, we tested for sequence conservation of cis-regulatory loci and gene exons in species with and without the 10 chromosome fusion events. Cis-regulatory elements were derived from ATAC-seq data for mid-pupal wing tissue from *H. erato* and *D. iulia* (15, 27) ATAC-seq data were processed as previously described (15) and peaks were called using F-seq (77). To reduce discrepancies between genome assembly annotations, which may have different degrees of gene fragmentation, we used annotated exons from *H. erato* and *D. iulia* to test for gene coding elements (27). A custom script was then used to perform a reciprocal best-hit BLAST search with an e-value threshold of e-10 on ATAC-seq peaks and gene exons to identify conserved functional DNA sequences. Peak and exon sets were split into files for each of the 30 (*D. iulia*) and 20 (*H. erato*) chromosomes. These datasets were then BLASTed against the *H. erato lativitta* (14), *H. melpomene* (78), *E. isabella, D. iulia, M. cinxia* (28), and *D. plexippus* (79) genome assemblies. Regression analysis and calculation of correlation coefficients was performed using the Python Statsmodels package.

## References

1. P. G. D. Feulner, R. De-Kayne, Genome evolution, structural rearrangements and speciation. J. Evol. Biol. 30, 1488–1490 (2017).

2. N. De Storme, A. Mason, Plant speciation through chromosome instability and ploidy change: Cellular mechanisms, molecular factors and evolutionary relevance.Curr. Plant Biol. 1, 10–33 (2014).

3. T. Dobzhansky, M. J. D. White, Animal Cytology and Evolution. Cambridge Univ. Press. 1983 (1977) https:/doi.org/10.2307/2405365.

4. H. C. Hauffe, J. B. Searle, Chromosomal heterozygosity and fertility in house mice (Mus musculus domesticus) from Northern Italy. Genetics (1998).

5. J. M. de Vos, H. Augustijnen, L. Bätscher, K. Lucek, Speciation through chromosomal fusion and fission in Lepidoptera. Philos. Trans. R. Soc. Lond. B. Biol. Sci. 375, 20190539 (2020).

6. R. F. Guerrero, M. Kirkpatrick, Local adaptation and the evolution of chromosome fusions. Evolution (N. Y). (2014) https:/doi.org/10.1111/evo.12481.

7. D. H. Lunt, S. Kumar, G. Koutsovoulos, M. L. Blaxter, The complex hybrid origins of the root knot nematodes revealed through comparative genomics. PeerJ 2014, 1–25 (2014).

8. H. Fradin, et al., Genome Architecture and Evolution of a Unichromosomal Asexual Nematode. Curr. Biol. 27, 2928–2939.e6 (2017).

9. B. Charlesworth, Mutation-selection balance and the evolutionary advantage of sex and recombination. Genet. Res. (Camb). (1990) https:/doi.org/10.1017/S0016672308009658.

10. A. D. Cutter, B. A. Payseur, Genomic signatures of selection at linked sites: Unifying the disparity among species. Nat. Rev. Genet. (2013) https:/doi.org/10.1038/nrg3425.

11. R. B. Corbett-Detig, D. L. Hartl, T. B. Sackton, Natural Selection Constrains Neutral Diversity across A Wide Range of Species. PLoS Biol. (2015) https:/doi.org/10.1371/journal.pbio.1002112.

12. A. Mackintosh, et al., The determinants of genetic diversity in butterflies. Nat. Commun. (2019) https:/doi.org/10.1038/s41467-019-11308-4.

13. D. Villar, et al., Enhancer evolution across 20 mammalian species. Cell 160, 554–566 (2015).

14. J. J. Lewis, K. R. L. van der Burg, A. Mazo-Vargas, R. D. Reed, ChIP-Seq-Annotated Heliconius erato Genome Highlights Patterns of cis-Regulatory Evolution in Lepidoptera. Cell Rep. 16, 2855–2863 (2016).

15. J. J. Lewis, R. D. Reed, Genome-Wide Regulatory Adaptation Shapes Population-Level Genomic Landscapes in Heliconius. Mol. Biol. Evol. (2019) https:/doi.org/10.1093/molbev/msy209.

16. S. Kim, et al., Comparison of carnivore, omnivore, and herbivore mammalian genomes with a new leopard assembly. Genome Biol. 17, 1–12 (2016).

17. C. Concha, et al., Interplay between Developmental Flexibility and Determinism in the Evolution of Mimetic Heliconius Wing Patterns. Curr. Biol. (2019) https:/doi.org/10.1016/j.cub.2019.10.010.

18. C. G. Danko, et al., Dynamic evolution of regulatory element ensembles in primate CD4+ T cells. Nat. Ecol. Evol. 2(2018).

19. M. Moest, et al., Selective sweeps on novel and introgressed variation shape mimicry loci in a butterfly adaptive radiation. PLoS Biol. (2020) https:/doi.org/10.1371/journal.pbio.3000597.

20. S. M. Van Belleghem, et al., Complex modular architecture around a simple toolkit of wing pattern genes. Nat. Ecol. Evol. (2017) https:/doi.org/10.1038/s41559-016-0052.

21. J. W. Davey, et al., Major Improvements to the Heliconius melpomene Genome Assembly Used to Confirm 10 Chromosome Fusion Events in 6 Million Years of Butterfly Evolution. G3 Genes, Genomes, Genet. 6, 695–708 (2016).

22. M. Mandrioli, G. C. Manicardi, Holocentric chromosomes. PLoS Genet. 16, e1008918 (2020).

23. S. H. Martin, et al., Natural selection and genetic diversity in the butterfly heliconius melpomene. Genetics (2016) https:/doi.org/10.1534/genetics.115.183285

24. S. H. Martin, J. W. Davey, C. Salazar, C. D. Jiggins, Recombination rate variation shapes barriers to introgression across butterfly genomes. PLoS Biol. (2019) https:/doi.org/10.1371/journal.pbio.2006288.

25. N. B. Edelman, et al., Genomic architecture and introgression shape a butterfly radiation. 599, 594–599 (2019).

26. K. M. Kozak, et al., Multilocus species trees show the recent adaptive radiation of the mimetic heliconius butterflies. Syst. Biol. 64, 505–524 (2015).

27. J. J. Lewis, et al., The *Dryas iulia* genome supports multiple gains of a W chromosome from a B chromosome in butterflies. submitted (2021).

28. V. Ahola, et al., The Glanville fritillary genome retains an ancient karyotype and reveals selective chromosomal fusions in Lepidoptera. Nat. Commun. 5, 1–9 (2014).

29. M. Grbić, et al., The genome of Tetranychus urticae reveals herbivorous pest adaptations. Nature 479, 487–492 (2011).

30. G. Bernardi, et al., The mosaic genome of warm-blooded vertebrates. Science (80-.). 228, 953–958 (1985).

31. Z. Qu, et al., Millipede genomes reveal unique adaptations during myriapod evolution. PLoS Biol. 18, 1–24 (2020).

32. J. Schmitz, J. Brosius, Exonization of transposed elements: A challenge and opportunity for evolution. Biochimie 93, 1928–1934 (2011).

33. J. L. Campos, B. Charlesworth, The effects on neutral variability of recurrent selective sweeps and background selection. Genetics 212, 287–303 (2019).

34. W. G. Hill, A. Robertson, The effect of linkage on limits to artificial selection. Genet. Res. (1966) https:/doi.org/10.1017/S0016672300010156.

35. N. H. Barton, Linkage and the limits to natural selection. Genetics (1995).

36. D. C. Presgraves, Recombination enhances protein adaptation in Drosophila melanogaster. Curr. Biol. (2005) https:/doi.org/10.1016/j.cub.2005.07.065.

37. T. F. C. MacKay, et al., The Drosophila melanogaster Genetic Reference Panel. Nature (2012) https:/doi.org/10.1038/nature10811.

38. P. R. Haddrill, D. L. Halligan, D. Tomaras, B. Charlesworth, Reduced efficacy of selection in regions of the Drosophila genome that lack crossing over. Genome Biol.(2007) https:/doi.org/10.1186/gb-2007-8-2-r18.

39. S. Koren, et al., Canu : scalable and accurate long- - - read assembly via adaptive k - - - mer weighting and repeat separation. 1–35 (2016).

40. M. Watson, A. Warr, Errors in long-read assemblies can critically affect protein prediction. Nat. Biotechnol. 37, 127–128 (2019).

41. K. M. Kozak, Macroevolution and phylogenomics in the adaptive radiation of Heliconiini butterflies (2015).

42. B. J. Walker, et al., Pilon: An integrated tool for comprehensive microbial variant detection and genome assembly improvement. PLoS One 9(2014).

43. M. J. Roach, S. A. Schmidt, A. R. Borneman, Purge Haplotigs: Synteny Reduction for Third-gen Diploid Genome Assemblies. bioRxiv, 286252 (2018).

44. B. H. Zhu, et al., P_RNA_scaffolder: A fast and accurate genome scaffolder using paired-end RNA-sequencing reads. BMC Genomics 19, 1–13 (2018).

45. M. Qin, et al., LRScaf: Improving Draft Genomes Using Long Noisy Reads. bioRxiv, 374868 (2018).

46. G. C. Xu, et al., LR-Gapcloser: A tiling path-based gap closer that uses long reads to complete genome assembly. Gigascience 8, 1–14 (2018).

47. C. Camacho, et al., BLAST command line applications user manual (2013).

48. H. Tang, et al., ALLMAPS: Robust scaffold ordering based on multiple maps. Genome Biol. 16, 1–15 (2015).

49. A. Dobin, et al., STAR: ultrafast universal RNA-seq aligner. Bioinformatics 29, 15–21 (2013).

50. M. A. Depristo, et al., A framework for variation discovery and genotyping using next-generation DNA sequencing data. Nat. Genet. (2011) https:/doi.org/10.1038/ng.806.

51. H. Li, et al., The Sequence Alignment/Map format and SAMtools. Bioinformatics 25, 2078–2079 (2009).

52. P. Rastas, Lep-MAP3: Robust linkage mapping even for low-coverage whole genome sequencing data. Bioinformatics 33, 3726–3732 (2017).

53. A. M. Bolger, M. Lohse, B. Usadel, Trimmomatic: A flexible trimmer for Illumina sequence data. Bioinformatics 30, 2114–2120 (2014).

54. K. J. Hoff, S. Lange, A. Lomsadze, M. Borodovsky, M. Stanke, BRAKER1: Unsupervised RNA-Seq-based genome annotation with GeneMark-ET and AUGUSTUS.Bioinformatics 32, 767–769 (2015).

55. K. J. Hoff, A. Lomsadze, M. Borodovsky, M. Stanke, “Whole-genome annotation with BRAKER” in Methods in Molecular Biology, (2019) https:/doi.org/10.1007/978-1-4939-9173-0_5.

56. A. Lomsadze, V. Ter-Hovhannisyan, Y. O. Chernoff, M. Borodovsky, Gene identification in novel eukaryotic genomes by self-training algorithm. Nucleic Acids Res.33, 6494–6506 (2005).

57. M. Stanke, M. Diekhans, R. Baertsch, D. Haussler, Using native and syntenically mapped cDNA alignments to improve de novo gene finding. Bioinformatics 24, 637–644 (2008).

58. A. Smit, R. Hubley, P. Green, RepeatMasker Open-4.0. 2013-2015. http://www.repeatmasker.org (2013).

59. G. S. C. Slater, E. Birney, Automated generation of heuristics for biological sequence comparison. BMC Bioinformatics 6, 1–11 (2005).

60. M. G. Grabherr, et al., Full-length transcriptome assembly from RNA-Seq data without a reference genome. Nat. Biotechnol. 29, 644–52 (2011).

61. H. Li, Minimap2: Pairwise alignment for nucleotide sequences. Bioinformatics 34, 3094–3100 (2018).

62. M. Pertea, et al., StringTie enables improved reconstruction of a transcriptome from RNA-seq reads. Nat. Biotechnol. 33, 290–295 (2015).

63. C. Trapnell, et al., Differential gene and transcript expression analysis of RNA-seq experiments with TopHat and Cufflinks. Nat. Protoc. 7, 562–578 (2012).

64. C. Trapnell, et al., Transcript assembly and quantification by RNA-Seq reveals unannotated transcripts and isoform switching during cell differentiation. Nat. Biotechnol. 28, 511–515 (2010).

65. D. Mapleson, L. Venturini, G. Kaithakottil, D. Swarbreck, Efficient and accurate detection of splice junctions from RNA-seq with Portcullis. Gigascience 7, 1–11 (2018).

66. A. Bateman, UniProt: A worldwide hub of protein knowledge. Nucleic Acids Res. (2019) https:/doi.org/10.1093/nar/gky1049.

67. G. M. Boratyn, et al., Domain enhanced lookup time accelerated BLAST. Biol. Direct 7, 12 (2012).

68. S. El-Gebali, et al., The Pfam protein families database in 2019. Nucleic Acids Res. (2019) https:/doi.org/10.1093/nar/gky995.

69. S. R. Eddy, Profile hidden Markov models. Bioinformatics 14, 755–763 (1998).

70. Y. J. Kang, et al., CPC2: A fast and accurate coding potential calculator based on sequence intrinsic features. Nucleic Acids Res. 45, W12–W16 (2017).

71. M. Krzywinski, et al., Circos: An information aesthetic for comparative genomics. Genome Res. (2009) https:/doi.org/10.1101/gr.092759.109.

72. D. I. Warton, R. A. Duursma, D. S. Falster, S. Taskinen, smatr 3-an R package for estimation and inference about allometric lines. Methods Ecol. Evol. (2012) https:/doi.org/10.1111/j.2041-210X.2011.00153.x.

73. D. J. Benjamin, et al., Redefine statistical significance. Nat. Hum. Behav. (2017) https:/doi.org/10.1038/s41562-017-0189-z.

74. A. R. Quinlan, I. M. Hall, The BEDTools manual. Genome (2010).

75. H. Li, Aligning sequence reads, clone sequences and assembly contigs with BWA-MEM. 00, 1–3 (2013).

76. H. Li, A statistical framework for SNP calling, mutation discovery, association mapping and population genetical parameter estimation from sequencing data.Bioinformatics (2011) https:/doi.org/10.1093/bioinformatics/btr509.

77. A. P. Boyle, J. Guinney, G. E. Crawford, T. S. Furey, F-Seq: A feature density estimator for high-throughput sequence tags. Bioinformatics (2008) https:/doi.org/10.1093/bioinformatics/btn480.

78. U. Kingdom, et al., Major Improvements to the Heliconius melpomene Genome Assembly Used to Confirm 10 Chromosome Fusion Events in 6 Million Years of Butterfly Evolution. 6, 695–708 (2016).

79. L. Gu, et al., Dichotomy of Dosage Compensation along the Neo Z Chromosome of the Monarch Butterfly. Curr. Biol. 29, 4071–4077.e3 (2019).

